# A conversion from slow to fast memory in response to passive motion

**DOI:** 10.1101/2021.03.09.434594

**Authors:** Mousa Javidialsaadi, Scott T. Albert, Jinsung Wang

**Author notes:** Correspondence: Jinsung Wang, 492 Enderis Hall, University of Wisconsin – Milwaukee, WI 53201, USA. **Data availability statement:** The data that support the findings of this study are available from the corresponding author upon reasonable request. The first and second authors contributed equally to this work.

## Abstract

When the same perturbation is experienced consecutively, learning is accelerated on the second attempt. This savings is a central property of sensorimotor adaptation. Current models suggest that these improvements in learning are due to changes in the brain’s sensitivity to error. Here, we tested whether these increases in error sensitivity could be facilitated by passive movement experiences. In each experimental group, a robot moved the arm passively in the direction that solved the upcoming rotation, with no visual feedback provided. Following that, participants adapted to a visuomotor rotation. Prior passive movements substantially improved motor learning, increasing total compensation in each group by approximately 30%. Similar to savings, a state-space model suggested that this improvement in learning was due to an increase in error sensitivity, but not memory retention. When we considered the possibility that learning was supported by parallel fast and slow adaptive processes, a striking pattern emerged: whereas initial improvements in learning were driven by a slower adaptive state, increases in error sensitivity gradually transferred to a faster learning system with the passage of time. These findings suggest that passive errors engage motor learning systems, but the resulting behavioral patterns migrate between slow and fast adaptive circuits as the passive memory is consolidated.

## Introduction

When we make a reaching movement, our brain predicts how our motor commands will change our sensory state. Perturbations to our movements create an error between the predicted sensory state and the observed sensory state (Popa & Ebner, 2019; Shadmehr et al., 2010). These sensory prediction errors cause learning, or changes to our motor commands on the next movement (Krakauer et al., 2000; Mazzoni & Krakauer, 2006). One hallmark of sensorimotor learning is savings: when people are exposed to the same perturbation twice in a row, they learn more rapidly during the second exposure (Smith et al., 2006; Kitago et al., 2013). Some models suggest that savings occurs because the brain recalls the movements it made in the past (Haith et al., 2015; Morehead et al., 2015). Other models suggest that savings occurs because the brain becomes more sensitive to errors it has experienced in the past (Gonzalez-Castro & Hadjiosif et al., 2014; Herzfeld et al., 2014; Albert et al., 2020). A common assumption across both models, however, is that savings is driven by active movement experiences.

But changes in learning rate can also occur even when people have not actively moved in the past. For example, recent studies have shown that passive sensorimotor experiences can improve active motor learning (Lei et al., 2016; Bao et al., 2017; Lei et al., 2017; Tays et al., 2020). Robot-assisted experiences enhance our ability to acquire novel motor skills (Reinkensmeyer and Patton, 2009; Bara and Gentaz, 2011; Basteris et al., 2012; Beets et al., 2012), facilitate motor recovery (Sakamoto et al., 2012; Tays et al., 2020), and improve motor function in hemiparetic stroke patients (Aisen et al., 1997; Krebs et al., 1998; Riener et al., 2005; Kahn et al., 2006; Vergaro et al., 2010).

For active movements, current state-space models (Smith et al., 2006; McDougle et al., 2015; Albert et al., 2018; Coltman et al., 2019; Orozco et al., 2020) posit that adaption is due to an interplay between at least two parallel adaptive processes: a slowly-adapting system (slow process) which responds weakly to error, but strongly retains its memory, and a rapidly-adapting system (fast process) which responds strongly to error, but weakly retains its memory. An increase in either system’s sensitivity to error is widely associated with savings in sensorimotor learning paradigms (Gonzalez-Castro et al., 2014; Herzfeld et al., 2014; Leow et al., 2016; Albert et al., 2020). Does passive training, then, enhance active motor learning by increasing error sensitivity in slow and/or fast learning processes?

Here we investigated this question in a visuomotor rotation experiment. In this paradigm, participants move a cursor towards a target, but the cursor’s position is rotated relative to the hand’s path. We used a robot to passively create a memory that could help solve a visuomotor rotation. Following this passive training, participants actively moved a rotated cursor to various visual targets. We varied the time between passive and active training to allow the passive memory to consolidate, and investigated how this consolidation altered the error sensitivities in the slow and fast adaptive processes (Shadmehr & Brashers-Krug, 1997; Caithness et al., 2004; Krakauer et al., 2005; Lerner & Albert et al., 2020).

## Materials & Methods

### Participants

Twenty-eight healthy volunteers (17 males, 11 females; aged: 18-35) participated in this study. All subjects were right-handed as assessed by the Edinburgh handedness inventory (Oldfield, 1971). Each subject signed a consent form that was approved by the University of Wisconsin-Milwaukee Institutional Review Board. Participants were randomly assigned to one of three experimental groups or a control group.

### Apparatus

Participants were seated in a robotic exoskeleton (KIMARM, BKIN Technologies Ltd, Kingston, ON, Canada) that provided gravitational support to both arms. The exoskeleton was positioned so that the participant’s arm was hidden underneath a horizontal display (Fig. 1A). To track the hand’s position, a small cursor was projected onto the display, over the participant’s index fingertip. During each trial, the KINARM projected visual stimuli (a start position and a target) onto the display, so that they appeared to be within the same plane as the arm. The visual stimuli consisted of a centrally located start circle (2 cm in diameter) and one of four target circles (2cm in diameter) located 10 cm away from the start target (Fig. 1B). We sampled the hand’s position in the x-y plane at 1000 Hz. Position data were low pass filtered at 15 Hz, and then differentiated to obtain velocity. Post-processing, analysis, and modeling were conducted in MATLAB R2018a (The MathWorks Inc., Natick, MA).

**Figure 1.**
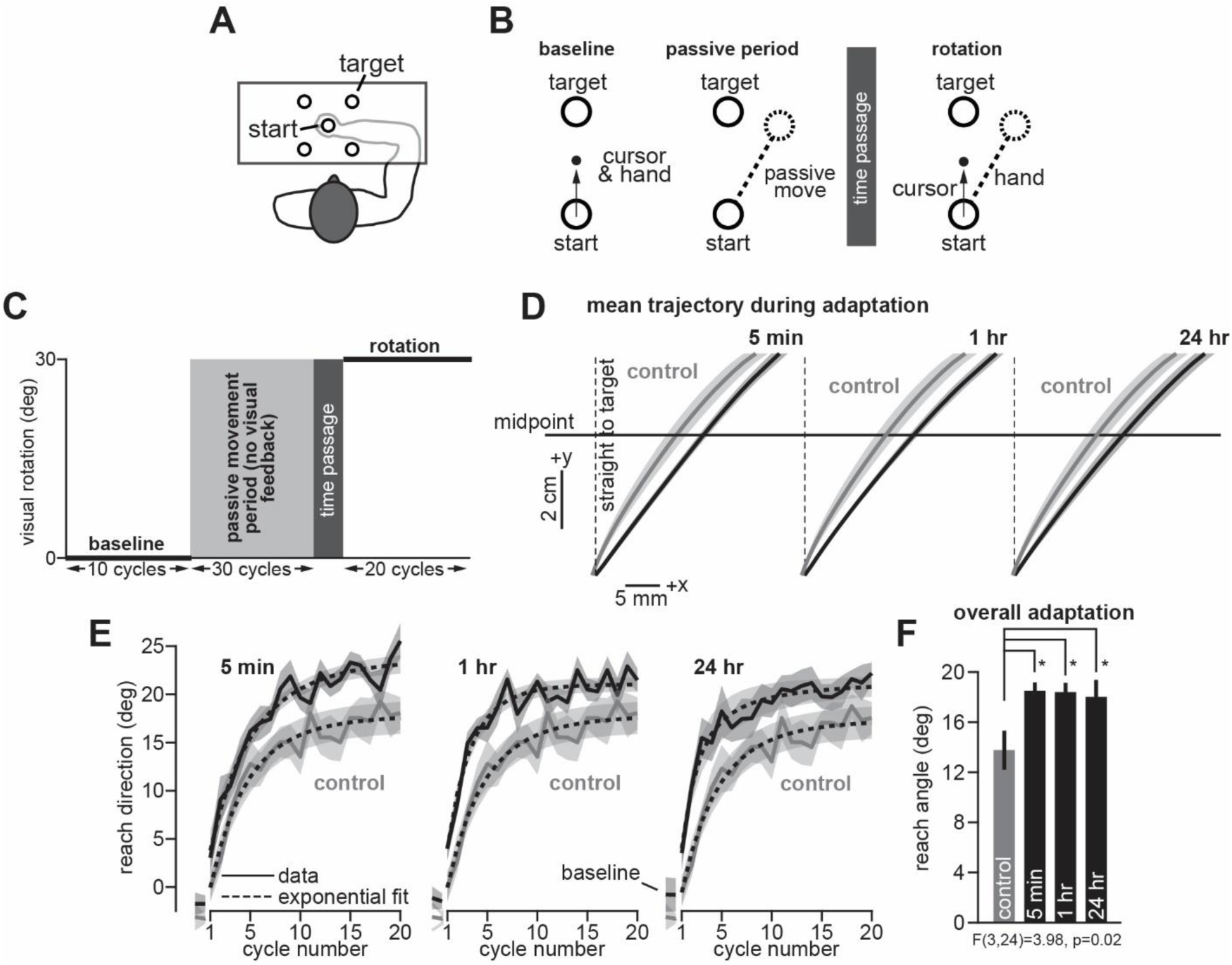
Passive movement experiences facilitate active motor adaptation. **A.** Participants were seated in a KINARM exoskeleton. The arm was placed underneath a horizontal display. **B.** The experiment began with a baseline phase, where the cursor moved with the hand. During the passive movement period, participants were shown a target in the 45°, 135°, 225° or 315° direction, but the KINARM passively moved the hand 30° CW relative to the visual target. No visual feedback was provided. Then participants took a break (time passage). When participants returned to the task, they started an active rotation period where the cursor was rotated 30° to the hand. Thus, to “solve” the rotation, participants needed to reach along the same path traversed during the passive period. **C.** Complete experiment paradigm. Experimental groups were divided into a 5 min, 1 hr, or 24 hr time passage condition. **D.** Average reach trajectories over the 20 rotation cycles. The control group (no passive training) is shown in gray. Reach angles for 5 min group (left), 1 hr group (middle), and 24 hr group (right) are shown in black. We calculated the reach angle between the hand and target cursor, at the movement’s midpoint (5 cm displacement). **E.** Cycle-by-cycle hand angles at reach midpoint. **F.** We calculated the mean reach angle across the rotation period. Statistics show post-hoc Tukey’s test following one-way ANOVA (* indicates p<0.05). Error bars show mean ± SEM.

### Experimental Design

Participants were assigned to one of three experimental groups (n=7/group). Each experimental group experienced four separate periods: (1) a baseline reaching period, (2) a passive movement period, (3) a break in time, and (4) a visuomotor rotation period (Fig. 1C). To determine how passive movements altered reaching behavior, we compared behavior in the experimental groups to that of a control group (n=7) that did not experience the passive movement period prior to rotation learning.

In the baseline period (Fig. 1B, baseline), participants moved their arm to each target over a 10-cycle reaching period. On each trial, continuous visual feedback was provided via a cursor over the index fingertip. Participants were instructed to move rapidly and accurately to the target location. The trial ended 1.5 sec after the target was presented. To begin the next trial, the participant had to move their hand back to the central start position.

In the passive movement period, the KINARM moved the participant’s right arm along a straight 10 cm minimum jerk trajectory. On each trial, a visual target was displayed, but the robot passively moved the arm along a path that was rotated 30° clockwise (CW) to the visual target (Fig. 1B, passive period). Critically, this CW rotation served as the “solution” to a counterclockwise (CCW) visuomotor rotation that participants had not yet experienced. Participants were told to relax their arm and not resist the passive motion. The passive period consisted of 30 cycles (4 trials in a cycle, 1 to each target in a pseudorandom order). No visual feedback was provided on these trials. Thus, participants received only proprioceptive information about the error between their arm’s path and the visual target. The passive movement period terminated with a time-delay which distinguished each experimental group: 5-min, 1-hr, or 24-hr delay. Once the delay concluded, the rotation period began.

The experiment ended with an active visuomotor rotation period that lasted 20 cycles (80 trials total). On rotation trials, the cursor was rotated 30° CCW relative to the hand path (Fig. 1B, rotation). As noted above, the “solution” to this rotation would be to counter-rotate the reach path by 30° CW relative to the target. Critically, this CW rotation would coincide with the movement path experienced during the passive period. Thus, participants in each experimental had been given the proprioceptive experience of this “solution”, though this was never explicitly revealed to each participant. Participants in the control group, however, had never experienced the passive movement period. Thus, there was no prior memory to draw upon during the rotation period.

### Empirical Data Analysis

Data analysis was performed using MATLAB (Mathworks, Natick, MA). The pointing angle at the reaching movement’s midpoint (5 cm displacement) was used as our performance measure, calculated as the hand’s angular position relative to the line segment connecting the start and target positions. Data were then averaged within each 4-trial cycle.

Here we considered how passive training altered adaptation during the rotation period. Fig. 1D illustrates the average reach trajectory over the 20-cycle rotation period. To calculate these mean hand paths, we artificially rotated each reach trajectory to create a 90° angle between the start location and the target (i.e., movement directed along the y-axis alone). Next, we interpolated the hand’s lateral deviation (the x-axis in Fig. 1D) at 25 points evenly spaced between the start location and an 8 cm displacement towards the target (the y-axis in Fig. 1D). Fig. 1E shows hand angles on the last two baseline cycles, as well as the learning curve during the 20-cycle rotation period. To assess overall adaptation, we averaged each participant’s midpoint hand angle over the 20-cycle rotation period (Figs. 1F).

To aid visual inspection, we fit an exponential model to the mean learning curve in each group (Fig. 1E, dashed line, exponential fit). This exponential curve tracked how reach angle (*y*) changed over each cycle (*t*, starting at 0):

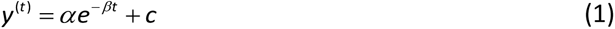

Here *α* and *c* determine the initial reach angle and asymptotic reach angle. The parameter β represents the participant’s learning rate during the adaptation period. We fit this exponential function to each group’s mean data in the least-squares sense using *fmincon* in MATLAB R2018a. We repeated the fitting procedure 20 times, each time varying the initial parameter guess that seeded the algorithm. We selected the model parameters which minimized squared error over all 20 repetitions.

We observed that participants who completed passive training exhibited improved adaptation (Fig. 1F). These improvements could have been due to 3 non-exclusive causes. First, the differences in adapted reach angles may have been due to biases in reach angles that lingered from the baseline reach period (Hypothesis 1). Second, improvements in adaptation may have been due to use-dependent biases in reach angles induced by passive training (Hypothesis 2). Third, improved adaptation may have been due to an enhancement in trial-to-trial learning induced by passive training (Hypothesis 3). To evaluate each hypothesis, we analyzed separate reach periods during the experiment protocol.

For Hypothesis 1 (lingering baseline biases), we calculated the reach angles at the very end of the baseline period in each participant. We calculated these angles both on the last 2 baseline trials (Figs. 2A-C) and the last 4 baseline trials. For Hypothesis 2 (use-dependent bias at the start of the rotation period), we calculated the reach angles at the very start of the rotation period in each participant (Figs. 2D-F). We calculated these angles both on the initial 2 rotation trials and the initial 4 rotation trials.

**Figure 2.**
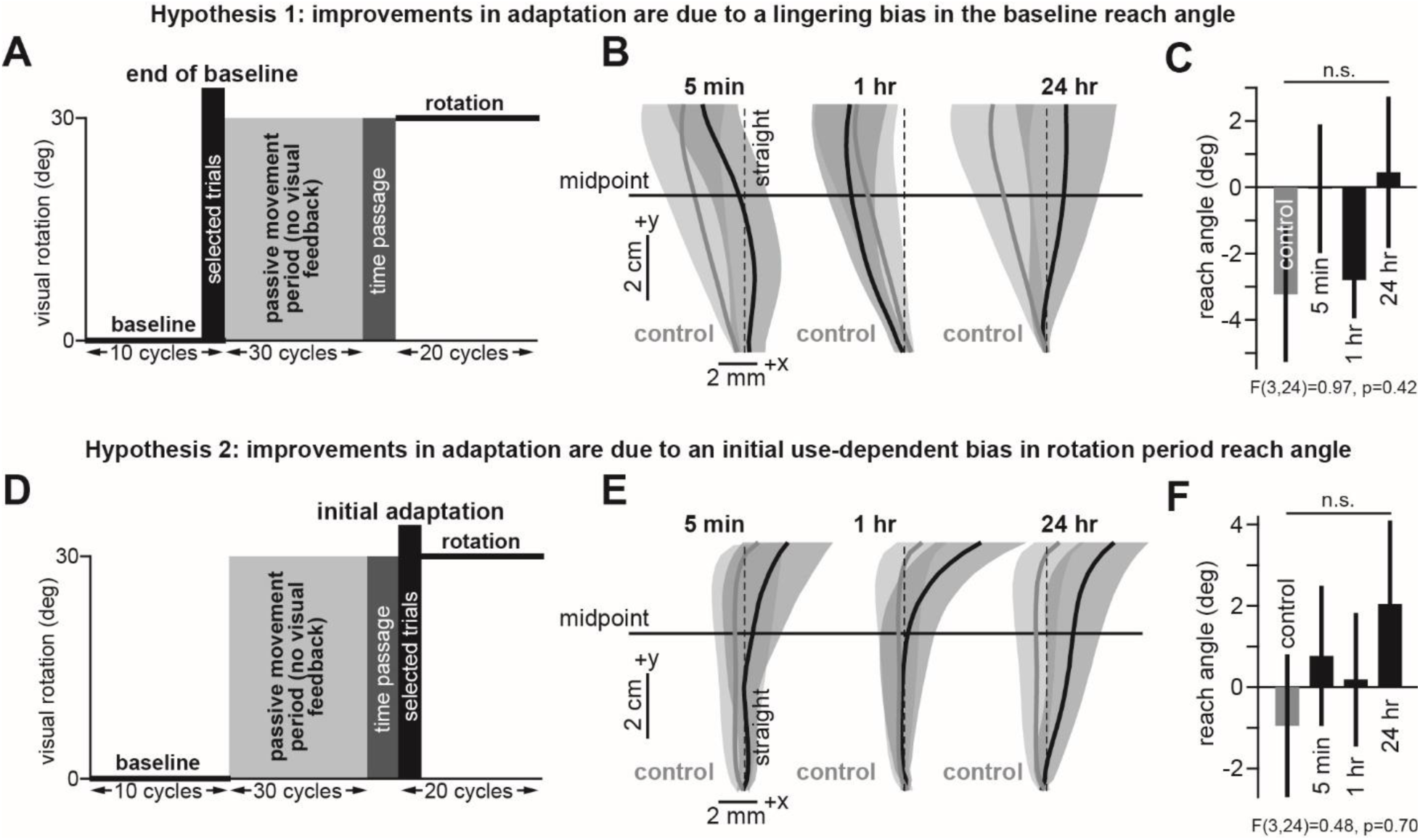
Improvements in adaptation were unrelated to biases in reach angle. **A.** We measured reach angles at the very end of the baseline period (end of baseline, last 2 baseline trials). **B.** Average trajectory on the end of baseline trials. **C.** Mean reach angle over the end of baseline trials. **D.** We measured reach angles at the start of the rotation period (initial adaptation, first 2 rotation trials). **E.** Average trajectory on the initial adaptation trials. **F.** Mean reach angle over the initial adaptation trials. Statistics indicate the group main effect in a one-way ANOVA (n.s. denotes p>0.05). Error bars show mean ± SEM.

For Hypothesis 3 (passive training improves learning), we noted that adaptation is supported by slow-learning processes and fast-learning processes (Smith et al., 2006). Fast processes are dominant early during adaptation. Slow processes are dominant late in adaptation. Thus, we measured adaptation both early and late during the learning process. To determine the point at which fast adaptation switched to slow adaptation, we calculated a cycle-by-cycle learning rate measure, i.e., the adaptation curve’s slope on each cycle. To do this, we used a second-order finite difference (Fig. 3A) to approximate the derivative of the learning curves in Fig. 1E on each cycle. We observed that adaptation exhibited rapid learning on adaptation cycles 1 and 2, intermediate learning on cycle 3, and slow learning on cycle 4 and beyond. Thus, we divided the adaptation period into a rapid early learning period (Figs. 3B&C; cycles 1 and 2), as well as a slow late learning period (Figs. 3D&E; rotation cycles 3-20). We calculated the mean reach angle over these two periods in Figs. 3C&E. In addition, we calculated a normalized early learning metric by subtracting off the baseline reaching angle exhibited on the last baseline cycle.

**Figure 3.**
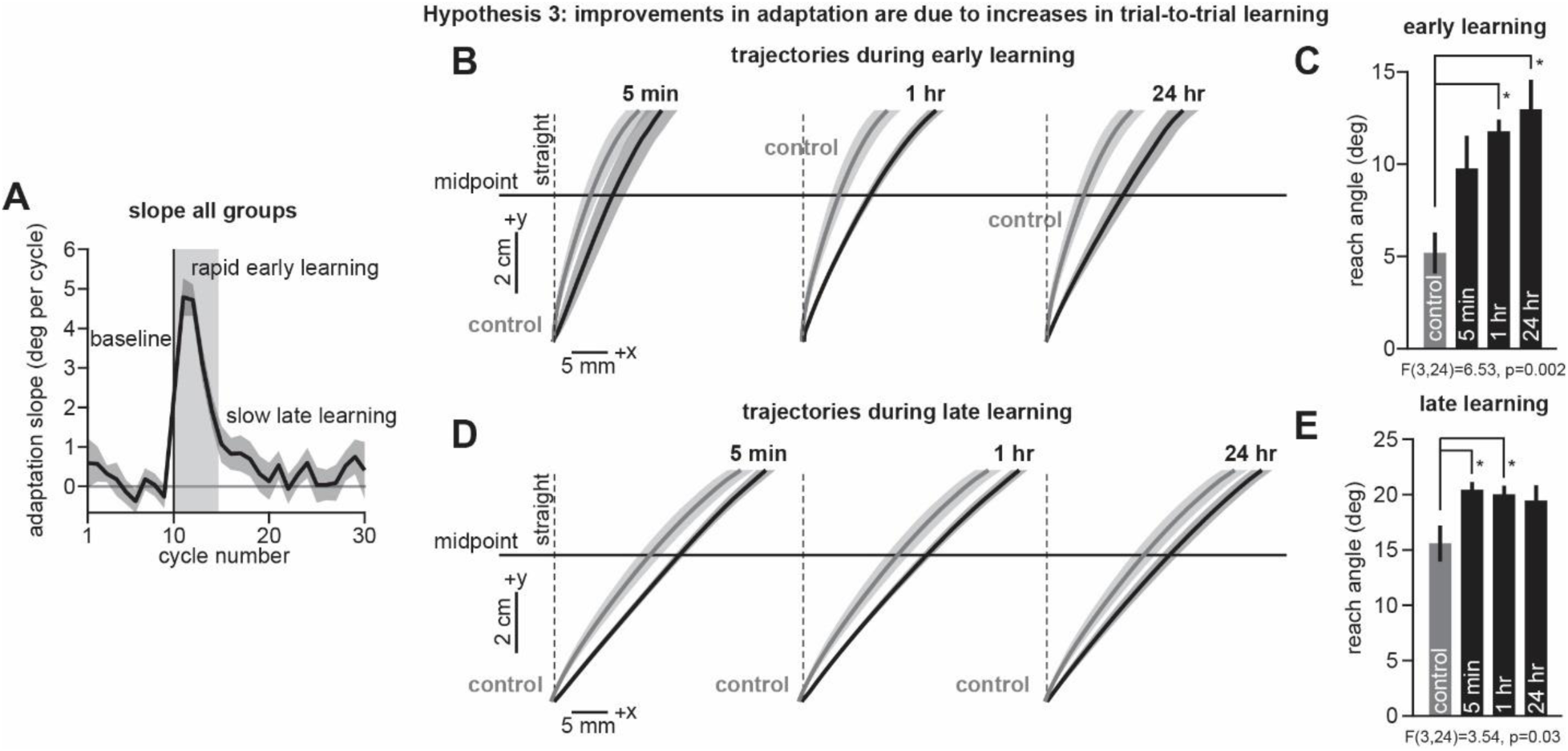
Passive training enhances early and late motor adaptation. **A.** We calculated an instantaneous adaptation rate measure on each cycle (using a second-order finite difference, Bozorgpour., 2023; Bozorgpour et al., 2023). Here data are averaged across all experimental groups. The adaptation rate was large during an initial 2-cycle period (rapid early learning). The adaptation rate was small late during adaptation (slow late learning). **B.** Mean reach trajectory over the rapid early learning period. Control group (no passive training) is shown in gray. Experimental groups (5 min, 1 hr, and 24 hr) are shown in black. **C.** Mean reach angle over the rapid early learning period. **D.** Mean reach trajectory over the slow late learning period. **E.** Mean reach angle over the late slow learning period. Statistics: in each panel, statistics show post-hoc Dunnett’s test against the control group following one-way ANOVA (*p<0.05). Error bars show mean ± SEM.

### State-space learning model: one-state

To better understand the adaptation process, we used a state-space model (Smith et al., 2006). The state-space model posits that learning consists of cycle-to-cycle error-based learning as well as cycle-by-cycle memory retention (i.e., forgetting). The forgetting process is controlled by a retention factor (*a*), which specifies how much adaptation is retained from one cycle to the next. The learning process is controlled by the participant’s error sensitivity (*b*), which specifies how much is learned from a given error (*e*). These processes together determine how the participant’s internal state (*x*) changes over time, in the presence of internal state noise (ε_x_, normal with zero mean, SD=σ_x_):

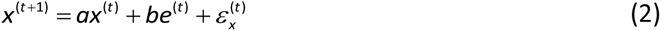

Eq. (2) allows us to ascribe any differences in performance during the adaptation period to meaningful quantities: retention (*a*) and error sensitivity (*b*).

Note, however, that the internal state (*x*) is not a measurable quantity. Rather, on each cycle, the motor output (reach angle) is measured. This reach angle (*y*) directly reflects the subject’s internal state, but is corrupted by execution noise (ε_x_, normal with zero mean, SD=σ_y_) according to:

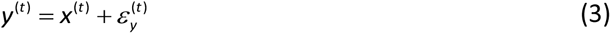

Thus, Eqs. (2) and (3) represent a single module state-space model. We fit this model to each participant’s reach angles during the adaptation period using the Expectation-Maximization (EM) algorithm (Albert & Shadmehr, 2018). EM is an algorithm that conducts maximum likelihood estimation in an iterative process. We used EM to identify the model parameters {*a*, *b*, *x_1_*, σ_x_, σ_y_, σ*_1_*} that maximized the likelihood of observing the data (note that *x_1_* and σ *_1_* represent the participant’s initial state and variance, respectively). We conducted 10 iterations, each time changing the initial parameter guess that seeded the algorithm. We selected the parameter set that maximized the likelihood function across all 10 iterations. We compared the retention factors and error sensitivities predicted by the model in Figs. 4B&C.

**Figure 4.**
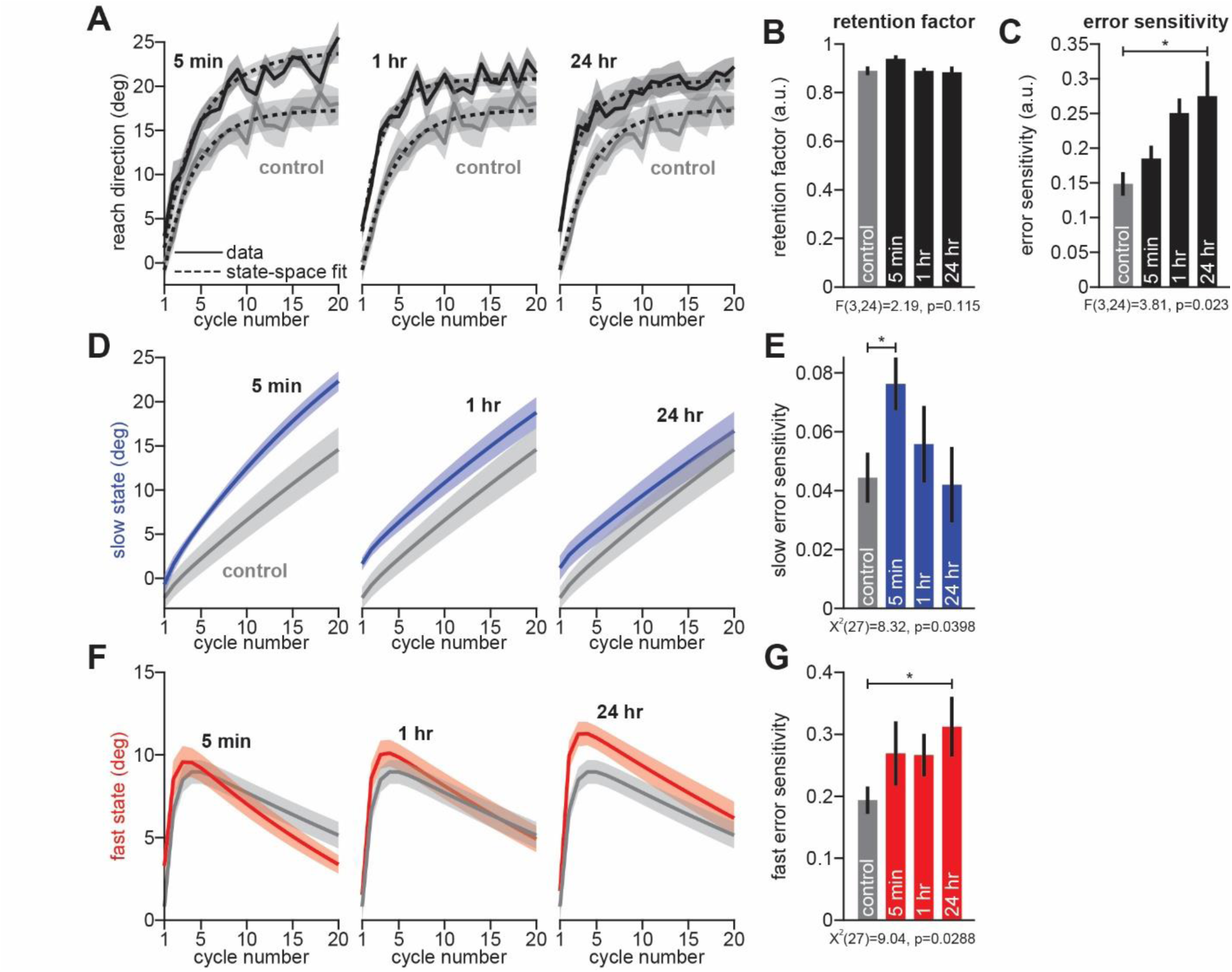
Passive training improves motor learning by increasing sensitivity to error. **A.** Here we show reach angles during the active rotation learning cycles. Control group (no passive training) is shown in gray. Experimental groups (5 min, 1 hr, and 24 hr) are shown in black. We fit the data with a single-module state-space model (dashed black line; mean across individual participants). The state-space model posited that adaptation was due to both error-based learning (controlled by error sensitivity) and trial-by-trial forgetting (controlled by retention factor). **B.** Here we show retention factor predicted by the single-module state-space model. **C.** Here we show error sensitivity predicted by the single-module state-space model. **D.** We fit a two-state model to individual participant behavior. Predicted slow state for experimental groups (5 min, 1 hr, and 24 hr) are shown in blue; time-delay between passive and active training increases left-to-right. Predicted slow state in control group is shown in gray. **E.** Here we provide slow state error sensitivity predicted by the two-state model. **F.** Same as in **D**, but for the fast state of learning. **G.** Same as in **E** but for the fast state error sensitivity. In each panel, statistics show post-hoc Tukey’s test (**C**) or Dunn’s test (**G**) following one-way ANOVA (**C**) or Kruskal-Wallis (**E** and **G**). Statistics: * indicates p<0.05, ** indicates p<0.01). Error bars show mean ± SEM.

### State-space learning model: two-state

The single-module state-space model describes learning as a single adaptive process. Motor adaptation, however, is believed to be comprised of multiple states, each with different timescales of learning. These states appear to be accurately summarized as a parallel two-state system, with adaptation supported by a slow adaptive process and a fast adaptive process (Smith et al., 2006; McDougle et al., 2015; Albert & Shadmehr, 2018; Coltman et al., 2019). The slow process exhibits slower error-based learning, but strong retention over time. The fast process exhibits faster error-based learning, but higher rates of forgetting. We wondered how these adaptive states are altered by consolidation of passive motor memory.

To answer this question, we fit a standard two-state model of learning to individual participant behavior. In this model, slow state and fast state adaptation are controlled by slow state and fast state retention factors (*a_s_* and *a_f_*, respectively), as well as slow state and fast state error sensitivities (*b_s_* and *b_f_*, respectively). As in Eq. (1), both internal states exhibit cycle-by-cycle learning and forgetting, in the presence of internal state noise (*ε*_x,s_ and *ξ*_x,s_, normal with zero mean, SD=*σ*_x_):

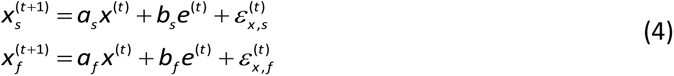

To enforce the traditional two-state dynamics (slow state has higher retention, fast state has higher error sensitivity), retention factors and error sensitivities in Eq. (4) are constrained such that *a_s_* > *a_f_* and *b_f_* > *b_s_*. As with the single-module state-space model, the two adaptive are not directly measurable. But together, they sum to produce the overall adapted reach angle, which is also corrupted by execution noise:

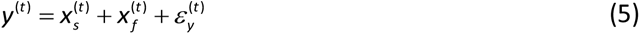

Together, Eqs. (4) and (5), along with the inequality constraints relating *a_s_*, *a_f_*, *b_s_*, and *b_f_*, constitute the two-state model of learning. The full parameter set consists of: {*a_s_*, *a_f_*, *b_s_*, *b_f_*, *x_s,1_*, *x_f,1_*, σ_x_, σ_y_, σ_1_}. Note that *x_s,1_*, and *x_f,1_* are the initial slow and fast state magnitudes, which are also estimated by the model.

We fit this model to individual participant data, using the EM algorithm (Albert & Shadmehr, 2018). Note that one’s ability to fit the two-state model greatly benefits from multiple trial conditions, such as set breaks, error-clamp periods, washout periods, and perturbation reversals (Albert & Shadmehr, 2018). However, the current experiment consisted of only perturbation trials with one orientation. Therefore, to improve our ability to robustly recover the two-state model parameters, we used a two-tiered fitting procedure (described below).

In our single-state model, we found that error sensitivity varied across experimental and control groups, but not retention. Therefore, to improve our ability to recover slow and fast error sensitivity (Albert & Shadmehr, 2018), we started by constraining the model’s retention factors (hypothesizing that these were not impacted by the passive movement period). To do this, we started by fitting the model to mean behavior in the 5 min, 1 hr, 24 hr, and control groups. Then we calculated the midpoint in the range spanned by slow and fast retention factors identified in each group. Lastly, we then fit the two-state model to individual participant behavior in each group, after fixing the slow and fast retention factors to these values.

When fitting the two-state model to either group behavior or individual participant behavior, we performed 10 iterations of the EM algorithm, each time varying the initial parameter guess that seeds the algorithm. We then selected the parameter set computed by EM with the greatest likelihood. We used these parameter sets to simulate the model’s slow and fast learning states in Figs. 4D&F. We compared the error sensitivities predicted by the model in Figs. 4E&G.

### Statistics

To compare the control and experimental groups, we conducted one-way ANOVAs in MATLAB R2018a. For post-hoc testing, we used Dunnett’s test to assess whether the behavior in the 5-min, 1-hr, and 24-hr delay groups differed from the control group. However, we used a non-parametric test to analyze our two-state model parameters, which are prone to outlying parameter estimates (Albert & Shadmehr, 2018; Coltman & Gribble, 2019). Thus, we used a Kruskal-Wallis test to evaluate differences in median error sensitivity (one outlier observed in the 1-hr slow state error sensitivity). For post-hoc testing following Kruskal-Wallis, we used Dunn’s test in IBM SPSS 25 (5-min, 1-hr, and 24-hr groups tested against control).

## Results

### Passive proprioceptive training facilitates active reach adaptation

The experiment began with a baseline period where participants reached to targets with veridical hand position feedback (Fig. 1B, baseline). Next, a passive training period ensued, where the robot moved the participant’s arm, without any visual feedback (Fig. 1B, passive period). On these trials, the robot moved the hand along a path rotated 30° CW relative to the target. Thus, each passive movement terminated in a proprioceptive “error” between the visual target and the hand. Following a break in time (Fig. 1B, time passage; 5 min, 1 hr, or 24 hr) participants returned to the task, but now actively moved to each target in the presence of a visuomotor rotation (Fig. 1B, rotation). How was adaptation altered by passive training?

To answer this question, we compared reaching movements during the rotation period (Fig. 1D, black) to those of a control group that never experienced passive training (Fig. 1D, gray). We calculated the angle of the hand’s path midway through the movement (Fig. 1D, midpoint). Hand angles rapidly adapted over the 20 rotation cycles (Fig. 1E) to reduce errors between the cursor and target. Remarkably, we observed that passive training greatly improved motor adaptation, increasing average compensation during the rotation period by 30% in all 3 experimental groups (Fig. 1F; one-way ANOVA, F(3,24)=3.98, p=0.02; post-hoc Dunnett’s test against control, 5 min: p=0.021, 1 hr: p=0.021, 24 hr: p=0.038).

It is possible that this improved adaptation was partially due to biases in reach angles that had developed during the baseline period. To assess whether baseline biases skewed our findings during the adaptation period (Hypothesis 1), we measured participant reach angles at the very end of the baseline period (Fig. 2A, end of baseline). At the end of baseline training, active reaching movements exhibited small biases ranging between about -3° and 0°. Critically, however, we did not detect any statistically significant differences in these initial biases between the experimental and control groups on the last two baseline trials (Fig. 2C; one-way ANOVA, F(3,24)=0.97, p=0.42), nor on the last four baseline trials (one-way ANOVA, F(3,24)=0.82, p=0.496). Thus, improvements in adaptation (Fig. 1F) were not due to any biases in reach angles that were carried over from the baseline period.

Instead, it was passive training that improved active motor adaptation. These improvements could have had two non-exclusive causes. First, passive training may have created a use-dependent bias in the initial reach angles during the adaptation period (Hypothesis 2). Second, passive training may have primed the motor learning system, increasing the amount participants adapted to visual errors (Hypothesis 3). Had passive training caused a use-dependent bias in reach angle (Hypothesis 2), this bias might have been present on the very initial rotation cycles (Fig. 2D, initial adaptation). However, when we assessed the initial reach angles during the rotation period (Figs. 2D&E), we did not detect any statistically significant differences across the experimental and control groups on the first two rotation trials (Fig. 2F; one-way ANOVA, F(3,24)=0.48, p=0.7), nor on the first four rotation trials (one-way ANVOA, F(3,24)=1.38, p=0.274).

Thus, improvements in adaptation were not observed in the initial reach angles during the rotation period; rather, they emerged as participants actively adapted to the rotation (Hypothesis 3). At what point did these changes emerge during rotation training? To investigate this, we first noted that adaptation is supported by multiple processes, some fast and others slow (Smith et al., 2006; Albert & Shadmehr, 2018), with fast processes dominant early during adaptation, and slow processes dominant late in adaptation. Indeed, trial-to-trial learning was very rapid over an initial 3-cycle period (Fig. 3A, gray region, rapid early learning), and then decreased abruptly over the remaining training epochs (Fig. 3A, slow late learning). Did passive training improve adaptation during the rapid initial learning phase, or during the late slow learning phase?

To answer this question, we compared each experimental group’s mean adaptation level over the early learning period with the control group (Fig. 3B). Interestingly, passive training strongly improved adaptation over the rapid early learning period (Fig. 3C; one-way ANOVA, F(3,24)=6.53, p=0.002), but the facilitation appeared stronger when more time had passed between passive and active training (post-hoc Dunnett’s test against control: 5 min: p=0.061, 1 hr: p=0.003, 24 hr, p=0.001). The same trend was observed when we normalized initial adaptation levels to each group’s baseline reach angles (one-way ANOVA, F(3,24)=3.2, p=0.041; post-hoc Dunnett’s test against control: 5 min: p=0.277, 1 hr: p=0.048, 24 hr, p=0.026). In sum, early learning was enhanced in the 1 hr and 24 hr groups.

Lastly, we measured each group’s mean adaptation over the late slowly adapting period (Fig. 3D). Here, we observed that less time passage between passive and active training produced stronger improvements in learning over the control group (Fig. 3E; one-way ANOVA, F(3,24)=3.54, p=0.03; post-hoc Dunnett’s test against control: 5 min: p=0.021, 1 hr: p=0.037, 24 hr: p=0.075).

In summary, we observed that passive training enhanced adaptation. These improvements were not due to a use-dependent bias in initial rotation reach angles. Rather, passive training appeared to improve active motor learning. But adaptation patterns varied with time passage between passive and active training. With short delays, improvements emerged later during rotation training. With long delays, improvements emerged earlier during rotation training.

### Passive movement experiences enhance sensitivity to error, but not retention

To better understand how passive training improved motor learning, we used a state-space model. In a state-space model, adaptation is an interplay between two processes: error-based learning and trial-by-trial forgetting. Error-based learning is controlled by the participant’s sensitivity to error (*b* in Eq. 2). Forgetting is controlled by the participant’s retention factor (*a* in Eq. 2). How did passive training alter each participant’s error sensitivity and retention?

To answer this question, we fit a single-module state-space model to individual participant reach angles in the experimental and control groups. The model (Fig. 4A, dashed lines) appeared to closely track each group’s learning curve. Next, we isolated the retention factor (Fig. 4B) and error sensitivity (Fig. 4C) predicted by the state-space model. Interestingly, passive training did not appear to alter the retention processes in any experimental group (Fig. 4B, one-way ANOVA, F(3,24)=2.19, p=0.115). On the other hand, passive training clearly impacted error sensitivity (Fig. 4C, one-way ANOVA, F(3,24)=3.81, p=0.023), which appeared to grow with time-delay duration; though only the 24 hr group showed a statistically significant difference relative to the control condition (post-hoc Dunnett’s tests against control, 5 min: p=0.72, 1 hr: p=0.061, 24 hr: p=0.017).

In sum, the state-space model predicted that passive training increased how sensitive participants were to errors experienced during the rotation period. But a puzzle remained: why is error sensitivity only increased in the 24 hr group (Fig. 4C), when all experimental groups showed a facilitation in learning (Fig. 1F)? We speculated that our state-space model was limited because it described adaptation as a single learning process. However, motor adaptation appears to be supported by at least two parallel adaptive processes: a slow state which has low error sensitivity but strong retention, and a fast state that has high error sensitivity but weak retention (Smith et al., 2006).

To better understand the error sensitivity patterns in Fig. 4C, we next applied a two-state learning model (Eqs. (4) & (5)) to the data. The two-state model predicted that adaptation was supported by the slow-adapting processes shown in Fig. 4D and the rapid-adapting processes shown in Fig. 4F. These states showed a striking trend: the slow-adapting process was more active with short delays between passive and active training (i.e., the 5 min group). The fast state of learning exhibited the opposite trend; it was more active with long delays between passive and active training (i.e., the 24 hr group).

These changes in the slow and fast states were due to a clear pattern in error sensitivity. The slow state’s error sensitivity increased nearly two-fold following the passive training period in the 5 min group (Fig. 4E, Kruskal-Wallis test, X^2^(27)=8.32, p=0.0398) but gradually returned to control levels as time elapsed between passive and active training (post-hoc Dunn’s tests against control, 5 min: p=0.024, 1 hr: p=1, 24 hr: p=1). On the other hand, the fast-state’s error sensitivity increased by 50% (Fig. 4G, Kruskal-Wallis test, X^2^(27)=9.04, p=0.0288) but only in the 24 hr group after the passage of time (post-hoc Dunn’s tests against control, 5 min: p=0.486, 1 hr: p=0.087, 24 hr: p=0.012). …

In sum, the two-state model indicated that passive training increased error sensitivity. Increases in error sensitivity improved visuomotor adaptation. With little time passage between passive and active training, error sensitivity was mostly improved in a slower adaptive state (Figs. 4D&E). With greater time passage, error sensitivity was mostly improved in a faster adaptive state (Figs. 4F&G).

## Discussion

When the same perturbation is experienced consecutively, learning is accelerated on the second attempt. This savings is a central property of sensorimotor adaptation that is observed across several motor effectors: the reaching system (Smith et al., 2006; Zarahn et al., 2008; Leow et al., 2013; Haith et al., 2015; Coltman et al., 2020; Yin & Wei, 2020), walking (Mawase et al., 2014; Day et al., 2018), and even the oculomotor system (Kojima et al., 2004). Current models have suggested that these improvements in learning are due to changes in the brain’s sensitivity to error (Gonzalez-Castro & Hadjiosif, 2014; Herzfeld et al., 2014; Mawase et al., 2014; Coltman et al., 2020). Here, we tested whether these increases in error sensitivity could be facilitated by passive experiences, as opposed to active movements.

Consistent with earlier work (Lei et al., 2016; Bao et al., 2017; Lei et al., 2017; Tays et al., 2020), we observed that passive training facilitated active motor adaptation. In each experimental group, a robot moved the arm passively in the direction that provided a solution to the upcoming rotation perturbation, with no visual feedback provided. This passive proprioceptive experience substantially altered subsequent motor learning, increasing total compensation in each group by approximately 30%. Similar to savings, the state-space model suggested that this improvement in learning was due to an increase in error sensitivity. Thus, passive memories appeared to increase the motor learning system’s sensitivity to error.

However, the state-space model’s prediction exhibited a puzzle: whereas all experimental groups showed improvements in total compensation, only the 24-hr group showed a statistically significant increase in sensitivity to error (Fig. 4C). This suggested that this model was missing a key component: multiple adaptive states. That is, converging evidence suggests that sensorimotor learning is supported by parallel processes (Smith et al., 2006; Mawase et al., 2014; McDougle et al., 2015; Coltman et al., 2020). The parallel adaptive states are well-approximated as a two-state system: one slow state that learns slowly but is decay-resistant, and one fast state that learns rapidly but also forgets rapidly. When we applied this two-state model to our data (Albert & Shadmehr, 2018), a new pattern emerged: passive training initially facilitated improvements in the slow state’s sensitivity to error (Fig. 4E). As time passed between passive and active training, these changes in the slow state’s error sensitivity dissipated; simultaneously, the fast process increased its error sensitivity (Fig. 4G).

How might passive training increase the brain’s sensitivity to error during active movements? Both passive movements (Carel et al., 2000) and active motor learning (Della-Maggiore & McIntosh, 2005; Herzfeld et al., 2015; Popa et al., 2012; Izawa et al., 2012; Tseng et al., 2007; Gibo et al., 2013; Maschke et al., 2004) are known to involve both the primary motor cortex (M1) and the cerebellum. One hypothesis is that prediction errors experienced during passive training engage Purkinje cells (P-cells) in the cerebellar cortex as well neurons in the deep cerebellar nuclei (Herzfeld et al., 2020; Lisberger et al., 1994; McCormick & Thompson, 1984; Perret et al., 1993). An interplay between P-cells and nuclear neurons may produce the changes in slow-state error sensitivity we observed early (5 min) after passive training. However, with time passage, plasticity in the cerebellum may be transferred to, and consolidated within M1 and S1 neurons via an unidentified mechanism (Kumar et al., 2019; Herzfeld et al., 2020; Orozco et al., 2020). These changes in the sensorimotor cortices may produce the increases in fast-state error sensitivity we observed with longer delays between passive and active training (24-hr group).

This first hypothesis draws upon the idea that passive training creates sensory prediction errors (SPEs). Current models have proposed that these SPEs engage the inferior olive, which in turn generate complex spikes in cerebellar P-cells (Herzfeld et al., 2015; Herzfeld et al., 2018). Associative learning paradigms such as eye-blink conditioning demonstrate that these SPEs and complex spikes are evoked even without active movement (Ohmae & Medina, 2015; Kim et al., 2020; Ito, 201; Sears & Steinmetz, 1991; Rasmussen et al, 2008). Thus, in our passive training paradigm, it is possible that the discrepancy between the visual target and the arm’s proprioceptive state (position) may have created an SPE and corresponding complex spikes in the cerebellar cortex (Popa & Ebner, 2019; Shadmehr et al., 2010). Repeated exposure to consistent prediction errors may have upregulated error sensitivity in the cerebellar cortex (Herzfeld et al., 2014; Albert et al., 2020), thus facilitating learning in the active condition. An alternate possibility is that passive training engaged the primary motor and somatosensory cortices, which then indirectly altered sensitivity to SPEs in the cerebellum via descending input. In either case, rTMS and tDCS over the cerebellum and M1 may provide a path to elucidating the various timescales of memory that are engaged by passive training.

A second hypothesis is that passive movements altered error sensitivity through use-dependent learning (Wang et al., 2015; Bao et al., 2017; Lei et al., 2017; Verstynen & Sabes, 2011; Lei et al., 2016; Diedrichsen et al., 2010; Jax & Rosenbaum, 2007; Scheidt et al., 2005; Diedrichsen, 2007), as a process by which motor plans are biased towards past movements. These biases arise in response to past active training (Verstynen & Sabes, 2011) as well as passive training (Diedrichsen et al., 2010). In our paradigm, it did not appear that passive training improved learning via a simple biasing in reach direction; when we measured participant reach angles on the initial rotation trials which followed passive training, we did not observe any biases in reach angle (Figs. 2D-F). Thus, while we have previously observed that passive training induces use-dependent biases in reach angle (Wang et al., 2015; Bao et al., 2017; Lei et al., 2017), it did not appear that these biases were strong enough to “pull” the arm toward the experienced movement direction in our experimental conditions. This may have been due to the uniform target arrangement in our experiment (Verstynen & Sabes, 2011) which could “average out” use-dependent learning across the 4 symmetric targets in the workspace.

A third intriguing hypothesis is that passive training engaged both a strategic explicit system that can be guided by instruction (Taylor et al., 2014; Mazzoni & Krakauer, 2006; McDougle & Taylor, 2019), and an implicit system that adapts without our conscious awareness (Avraham et al., 2020; Miyamoto et al., 2020; Javidialsaadi and Wang, 2020; Wang et al., 2019; Jo et al., 2022). Might an interplay between these two systems be related to the time-dependent error sensitivity patterns observed during rotation learning? Because we did not measure implicit or explicit learning, we cannot know the answer to this question. However, we did observe that reaction time (Fig. 5A) appeared elevated after passive training in the experimental groups, which commonly accompanies cognitive operations involved in explicit strategy use (Fernandez-Ruiz et al., 2011; Sakamoto and Kondo, 2012; Anguera et al., 2010; Georgopoulos and Massey, 1987; McDougle et al., 2019; Albert et al., 2020).

**Figure 5.**
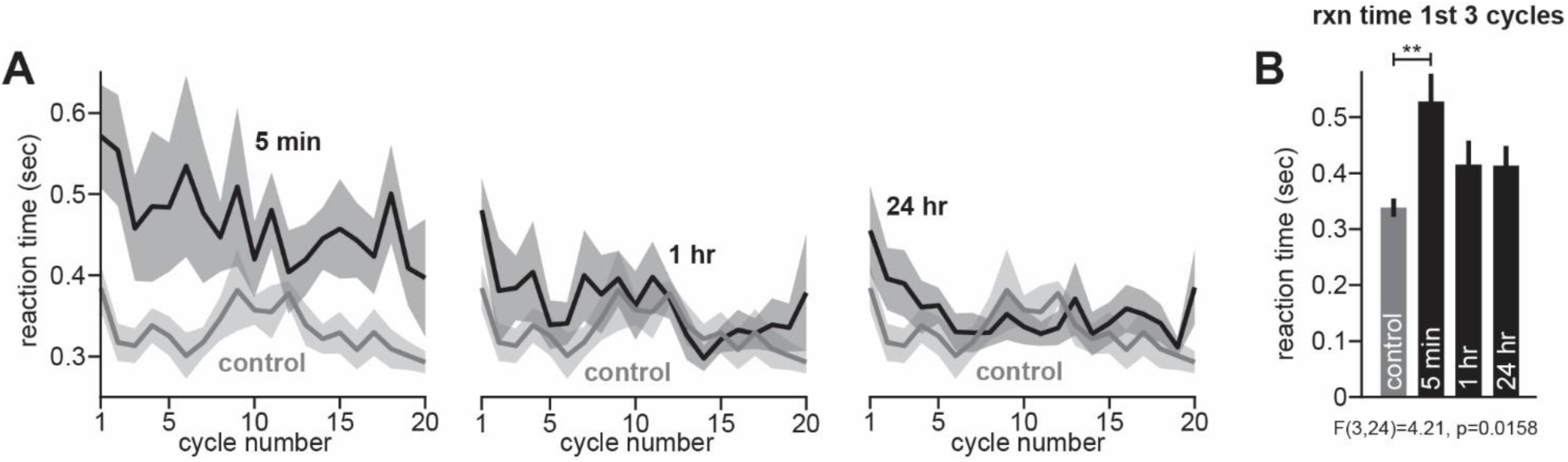
Changes in reaction time. **A.** We measured reaction time as the difference between target appearance and reach onset. Here we show reaction time during the perturbation period in experimental (black) and control (gray) groups. **B.** We investigated early differences in reaction time by calculating the mean reaction time over the first three cycles. Statistics show post-hoc Tukey’s test following one-way ANOVA (** indicates p<0.01). Error bars show mean ± SEM.

With that said, changes in reaction time were only statistically significant in the 5 min group (Fig. 5B) and appeared to dissipate quite rapidly in the 1-hr and 24-hr groups. If explicit strategies did contribute to increases in error sensitivity, it is also puzzling why reaction time was greatest in the 5 min group when this group was primarily enhanced by a slow learning state (Figs. 4D&E), which is thought to reflect implicit learning processes (McDougle et al., 2015). Nevertheless, the idea that passive training may evoke changes in both implicit and explicit processes is a fascinating possibility, which remains to be more formally tested.

## Conclusion

In summary, passive motor training led to improvements in active motor learning due to increases in error sensitivity. Fast and slow processes appeared to mediate these changes in learning but were each modulated over different timescales. With little time-passage between passive and active training, error sensitivity was enhanced in the slow adaptive state. With greater time-passage, error sensitivity was enhanced in the fast adaptive state. These changes in learning may have been mediated by interactions between the cerebral and cerebellar cortices, or potentially through implicit and explicit learning systems.

## Acknowledgements

This work was supported by a grant from the National Institutes of Health (F32NS095706). We dedicate this article to Dr. Fatemeh Pasand and Dr. Majid Chahardah Cheric for all their academic efforts and teachings. They passed away on 10 April 2018 and 26 June 2020, respectively.

## References

1. Albert, S. T., & Shadmehr, R. (2018). Estimating properties of the fast and slow adaptive processes during sensorimotor adaptation. Journal of neurophysiology, 119(4), 1367–1393.

2. Albert, Scott T., Jihoon Jang, Adrian M. Haith, Gonzalo Lerner, Valeria Della-Maggiore, John W. Krakauer, and Reza Shadmehr. “Competition between parallel sensorimotor learning systems.” bioRxiv (2020).

3. Albert, S. T. (2020). The role of error-based learning in movement and stillness (Doctoral dissertation, Johns Hopkins University).

4. Aisen, M. L., Krebs, H. I., Hogan, N., McDowell, F., & Volpe, B. T. (1997). The effect of robot-assisted therapy and rehabilitative training on motor recovery following stroke. Archives of neurology, 54(4), 443–446.

5. Anguera, J. A., Reuter-Lorenz, P. A., Willingham, D. T., & Seidler, R. D. (2010). Contributions of spatial working memory to visuomotor learning. Journal of cognitive neuroscience, 22(9), 1917–1930.

6. Avraham, G., Morehead, J. R., Kim, H. E., & Ivry, R. B. (2021). Reexposure to a sensorimotor perturbation produces opposite effects on explicit and implicit learning processes. PLoS biology, 19(3), e3001147.

7. Bao, S., Lei, Y., & Wang, J. (2017). Experiencing a reaching task passively with one arm while adapting to a visuomotor rotation with the other can lead to substantial transfer of motor learning across the arms. Neuroscience letters, 638, 109–113.

8. Bara, F., & Gentaz, E. (2011). Haptics in teaching handwriting: The role of perceptual and visuo-motor skills. Human movement science, 30(4), 745–759.

9. Basteris, A., Bracco, L., & Sanguineti, V. (2012). Robot-assisted intermanual transfer of handwriting skills. Human movement science, 31(5), 1175–1190.

10. Beets, I. A., Macé, M., Meesen, R. L., Cuypers, K., Levin, O., & Swinnen, S. P. (2012). Active versus passive training of a complex bimanual task: is prescriptive proprioceptive information sufficient for inducing motor learning?. PLoS One, 7(5), e37687.

11. Bozorgpour, R. (2023). Computational Explorations in Biomedicine: Unraveling Molecular Dynamics for Cancer, Drug Delivery, and Biomolecular Insights using LAMMPS Simulations. arXiv preprint arXiv:2311.13000.

12. Bozorgpour, R., Sheybanikashani, S., & Mohebi, M. (2023). Exploring the Role of Molecular Dynamics Simulations in Most Recent Cancer Research: Insights into Treatment Strategies. arXiv preprint arXiv:2310.19950.

13. Brashers-Krug, T., Shadmehr, R., & Bizzi, E. (1996). Consolidation in human motor memory. Nature, 382(6588), 252–255.

14. Carel, C., Loubinoux, I., Boulanouar, K., Manelfe, C., Rascol, O., Celsis, P., & Chollet, F. (2000). Neural substrate for the effects of passive training on sensorimotor cortical representation: a study with functional magnetic resonance imaging in healthy subjects. Journal of Cerebral Blood Flow & Metabolism, 20(3), 478–484.

15. Castro, L. N. G., Hadjiosif, A. M., Hemphill, M. A., & Smith, M. A. (2014). Environmental consistency determines the rate of motor adaptation. Current Biology, 24(10), 1050–1061.

16. Coltman, S. K., Cashaback, J. G., & Gribble, P. L. (2019). Both fast and slow learning processes contribute to savings following sensorimotor adaptation. Journal of neurophysiology, 121(4), 1575–1583.

17. Coltman, S. K., & Gribble, P. L. (2020). Time course of changes in the long-latency feedback response parallels the fast process of short-term motor adaptation. Journal of Neurophysiology, 124(2), 388–399.

18. Day, K. A., Leech, K. A., Roemmich, R. T., & Bastian, A. J. (2018). Accelerating locomotor savings in learning: compressing four training days to one. Journal of neurophysiology, 119(6), 2100–2113.

19. Della-Maggiore, V., & McIntosh, A. R. (2005). Time course of changes in brain activity and functional connectivity associated with long-term adaptation to a rotational transformation. Journal of neurophysiology, 93(4), 2254–2262.

20. Diedrichsen, J., Shadmehr, R., & Ivry, R. B. (2010). The coordination of movement: optimal feedback control and beyond. Trends in cognitive sciences, 14(1), 31–39.

21. Diedrichsen, J., Criscimagna-Hemminger, S. E., & Shadmehr, R. (2007). Dissociating timing and coordination as functions of the cerebellum. Journal of Neuroscience, 27(23), 6291–6301.

22. Fernandez-Ruiz, J., Wong, W., Armstrong, I. T., & Flanagan, J. R. (2011). Relation between reaction time and reach errors during visuomotor adaptation. Behavioural brain research, 219(1), 8–14.

23. Georgopoulos, A. P., & Massey, J. T. (1987). Cognitive spatial-motor processes. Experimental brain research, 65(2), 361–370.

24. Gibo, T. L., Criscimagna-Hemminger, S. E., Okamura, A. M., & Bastian, A. J. (2013). Cerebellar motor learning: are environment dynamics more important than error size?. Journal of neurophysiology, 110(2), 322–333.

25. Haith AM, Huberdeau DM, Krakauer JW. The influence of movement preparation time on the expression of visuomotor learning and savings. J Neurosci 35: 5109–5117, 2015.

26. Herzfeld, D. J., Hall, N. J., Tringides, M., & Lisberger, S. G. (2020). Principles of operation of a cerebellar learning circuit. Elife, 9, e55217.

27. Herzfeld, D. J., Kojima, Y., Soetedjo, R., & Shadmehr, R. (2018). Encoding of error and learning to correct that error by the Purkinje cells of the cerebellum. Nature neuroscience, 21(5), 736–743.

28. Herzfeld DJ, Vaswani PA, Marko MK, Shadmehr R. A memory of errors in sensorimotor learning. Science 345: 1349–1353, 2014.

29. Herzfeld, D. J., Kojima, Y., Soetedjo, R., & Shadmehr, R. (2015). Encoding of action by the Purkinje cells of the cerebellum. Nature, 526(7573), 439–442.

30. Huberdeau, D. M., Haith, A. M., & Krakauer, J. W. (2015). Formation of a long-term memory for visuomotor adaptation following only a few trials of practice. Journal of neurophysiology, 114(2), 969–977.

31. Huberdeau, D. M., Krakauer, J. W., & Haith, A. M. (2019). Practice induces a qualitative change in the memory representation for visuomotor learning. Journal of neurophysiology, 122(3), 1050–1059.

32. Ito, K., Doi, M., Kondo, T.: Feedforward Adaptation to a Varying Dynamical Environment during Reaching Movements, Journal of Robotics Mechatronics, 19, 4, 474–481 (2007).

33. Izawa, J., Criscimagna-Hemminger, S. E., & Shadmehr, R. (2012). Cerebellar contributions to reach adaptation and learning sensory consequences of action. Journal of Neuroscience, 32(12), 4230–4239.

34. Javidialsaadi, M., & Wang, J. (2021). Lack of interlimb transfer following visuomotor adaptation in a person with congenital mirror movements despite the awareness of the visuomotor perturbation. Brain and Cognition, 147, 105653.

35. Jax, S. A., & Rosenbaum, D. A. (2007). Hand path priming in manual obstacle avoidance: evidence that the dorsal stream does not only control visually guided actions in real time. Journal of Experimental Psychology: Human Perception and Performance, 33(2), 425.

36. Jo, Y., Javidialsaadi, M., & Wang, J. (2022). Facilitative effects of use-dependent learning on interlimb transfer of visuomotor adaptation in a person with congenital mirror movements. Human Movement Science, 84, 102973.

37. Kahn, L. E., Zygman, M. L., Rymer, W. Z., & Reinkensmeyer, D. J. (2006). Robot-assisted reaching exercise promotes arm movement recovery in chronic hemiparetic stroke: a randomized controlled pilot study. Journal of neuroengineering and rehabilitation, 3(1), 1–13.

38. Kim, H. E., Avraham, G., & Ivry, R. B. (2020). The psychology of reaching: action selection, movement implementation, and sensorimotor learning. Annual review of psychology, 72.

39. Kitago T, Ryan SL, Mazzoni P, Krakauer JW, Haith AM. Unlearning versus savings in visuomotor adaptation: comparing effects of washout, passage of time, and removal of errors on motor memory. Front Hum Neurosci 7: 307, 2013.

40. Kojima Y, Iwamoto Y, Yoshida K. Memory of learning facilitates saccadic adaptation in the monkey. J Neurosci 24: 7531–7539, 2004.

41. Krebs, H. I., Hogan, N., Aisen, M. L., & Volpe, B. T. (1998). Robot-aided neurorehabilitation. IEEE transactions on rehabilitation engineering, 6(1), 75–87.

42. Kumar, N., Manning, T. F., Ostry, D. G. (2019). Somatosensory cortex participates in the consolidation of human motor memory. PLoS Biol 17(10): e3000469.

43. Lei, Y., Bao, S., & Wang, J. (2016). The combined effects of action observation and passive proprioceptive training on adaptive motor learning. Neuroscience, 331, 91–98.

44. Lei, Y., Bao, S., Perez, M. A., & Wang, J. (2017). Enhancing generalization of visuomotor adaptation by inducing use-dependent learning. Neuroscience, 366, 184–195.

45. Leow LA, de Rugy A, Marinovic W, Riek S, Carroll TJ. Savings for visuomotor adaptation require prior history of error, not prior repetition of successful actions. J Neurophysiol 116: 1603–1614, 2016.

46. Leow, L. A., De Rugy, A., Loftus, A. M., & Hammond, G. (2013). Different mechanisms contributing to savings and anterograde interference are impaired in Parkinson’s disease. Frontiers in human neuroscience, 7, 55.

47. Lerner, G., Albert, S., Caffaro, P. A., Villalta, J. I., Jacobacci, F., Shadmehr, R., & Della-Maggiore, V. (2020). The origins of anterograde interference in visuomotor adaptation. Cerebral Cortex, 30(7), 4000–4010.

48. Lisberger, S. G. (1994). Neural basis for motor learning in the vestibuloocular reflex of primates. III. Computational and behavioral analysis of the sites of learning. Journal of Neurophysiology, 72(2), 974–998.

49. Maschke, M., Gomez, C. M., Ebner, T. J., & Konczak, J. (2004). Hereditary cerebellar ataxia progressively impairs force adaptation during goal-directed arm movements. Journal of neurophysiology, 91(1), 230–238.

50. Mawase, F., Shmuelof, L., Bar-Haim, S., & Karniel, A. (2014). Savings in locomotor adaptation explained by changes in learning parameters following initial adaptation. Journal of neurophysiology, 111(7), 1444–1454.

51. Mazzoni, P., & Krakauer, J. W., (2006). An implicit plan overrides an explicit strategy during visuomotor adaptation. Journal of neuroscience, 26(14), 3642–3645.

52. McCormick, D. A., & Thompson, R. F. (1984). Neuronal responses of the rabbit cerebellum during acquisition and performance of a classically conditioned nictitating membrane-eyelid response. Journal of Neuroscience, 4(11), 2811–2822.

53. McDougle, S. D., Bond, K. M., & Taylor, J. A., (2015). Explicit and implicit processes constitute the fast and slow processes of sensorimotor learning. Journal of Neuroscience, 35(26), 9568–9579.

54. McDougle, S. D., & Taylor, J. A. (2019). Dissociable cognitive strategies for sensorimotor learning. Nature communications, 10(1), 1–13.

55. Miyamoto, Y. R., Wang, S., & Smith, M. A. (2020). Implicit adaptation compensates for erratic explicit strategy in human motor learning. Nature neuroscience, 23(3), 443–455.

56. Morehead, J. Ryan, Salman E. Qasim, Matthew J. Crossley, and Richard Ivry. “Savings upon re-aiming in visuomotor adaptation.” Journal of neuroscience 35, no. 42 (2015): 14386–14396.

57. Ohmae, S., & Medina, J. F. (2015). Climbing fibers encode a temporal-difference prediction error during cerebellar learning in mice. Nature neuroscience, 18(12), 1798–1803.

58. Orozco, S. P., Albert, S. T., & Shadmehr, R. (2020). Spontaneous recovery and the multiple timescales of human motor memory. bioRxiv.

59. Perrett, S. P., Ruiz, B. P., & Mauk, M. D. (1993). Cerebellar cortex lesions disrupt learning-dependent timing of conditioned eyelid responses. Journal of Neuroscience, 13(4), 1708–1718.

60. Popa, L. S., & Ebner, T. J. (2019). Cerebellum, predictions and errors. Frontiers in cellular neuroscience, 12, 524.

61. Popa, L. S., Hewitt, A. L., & Ebner, T. J. (2012). Predictive and feedback performance errors are signaled in the simple spike discharge of individual Purkinje cells. Journal of Neuroscience, 32(44), 15345–15358.

62. Reinkensmeyer, D. J., & Patton, J. L. (2009). Can robots help the learning of skilled actions?. Exercise and sport sciences reviews, 37(1), 43.

63. Riener, R., Nef, T., & Colombo, G. (2005). Robot-aided neurorehabilitation of the upper extremities. Medical and biological engineering and computing, 43(1), 2–10.

64. Rasmussen, A., Jirenhed, D. A., & Hesslow, G. (2008). Simple and complex spike firing patterns in Purkinje cells during classical conditioning. The Cerebellum, 7(4), 563.

65. Sakamoto, T., & Kondo, T. (2012, November). Can passive arm movement affect adaptation to visuomotor rotation?. In 2012 IEEE International Conference on Development and Learning and Epigenetic Robotics (ICDL) (pp. 1–6). IEEE.

66. Shadmehr, R., Smith, M. A., & Krakauer, J. W. (2010). Error correction, sensory prediction, and adaptation in motor control. Annual review of neuroscience, 33, 89–108.

67. Scheidt, R. A., Conditt, M. A., Secco, E. L., & Mussa-Ivaldi, F. A. (2005). Interaction of visual and proprioceptive feedback during adaptation of human reaching movements. Journal of neurophysiology, 93(6), 3200–3213.

68. Sing, G. C., & Smith, M. A. (2010). Reduction in learning rates associated with anterograde interference results from interactions between different timescales in motor adaptation. PLoS Comput Biol, 6(8), e1000893.

69. Smith MA, Ghazizadeh A, Shadmehr R. Interacting adaptive processes with different timescales underlie short-term motor learning. PLoS Biol 4: e179, 2006.

70. Steinmetz, J. E., Sears, L. L., Gabriel, M., Kubota, Y., & Poremba, A. (1991). Cerebellar interpositus nucleus lesions disrupt classical nictitating membrane conditioning but not discriminative avoidance learning in rabbits. Behavioural brain research, 45(1), 71–80.

71. Taylor, J. A., Krakauer, J. W., & Ivry, R. B., (2014). Explicit and implicit contributions to learning in a sensorimotor adaptation task. Journal of Neuroscience, 34(8), 3023–3032.

72. Tays, G., Bao, S., Javidialsaadi, M., & Wang, J. (2020). Consolidation of use-dependent motor memories induced by passive movement training. Neuroscience Letters, 732, 135080.

73. Tong, C., & Flanagan, J. R. (2003). Task-specific internal models for kinematic transformations. Journal of Neurophysiology, 90(2), 578–585.

74. Tseng, Y. W., Diedrichsen, J., Krakauer, J. W., Shadmehr, R., & Bastian, A. J. (2007). Sensory prediction errors drive cerebellum-dependent adaptation of reaching. Journal of neurophysiology, 98(1), 54–62.

75. Vergaro, E., Casadio, M., Squeri, V., Giannoni, P., Morasso, P., & Sanguineti, V. (2010). Self-adaptive robot training of stroke survivors for continuous tracking movements. Journal of neuroengineering and rehabilitation, 7(1), 1–12.

76. Verstynen, T., & Sabes, P. N. (2011). How each movement changes the next: an experimental and theoretical study of fast adaptive priors in reaching. Journal of Neuroscience, 31(27), 10050–10059.

77. Wang, J., & Lei, Y. (2015). Direct-effects and after-effects of visuomotor adaptation with one arm on subsequent performance with the other arm. Journal of neurophysiology, 114(1), 468–473.

78. Wang, J., Bao, S., & Tays, G. D. (2019). Lack of generalization between explicit and implicit visuomotor learning. PloS one, 14(10), e0224099.

79. Wigmore, V., Tong, C., & Flanagan, J. R. (2002). Visuomotor rotations of varying size and direction compete for a single internal model in a motor working memory. Journal of Experimental Psychology: Human Perception and Performance, 28(2), 447.

80. Yin, C., & Wei, K. (2020). Savings in sensorimotor adaptation without an explicit strategy. Journal of neurophysiology, 123(3), 1180–1192.

81. Zarahn, E., Weston, G. D., Liang, J., Mazzoni, P., & Krakauer, J. W. (2008). Explaining savings for visuomotor adaptation: linear time-invariant state-space models are not sufficient. Journal of neurophysiology, 100(5), 2537–2548.

